# cDeepbind: A context sensitive deep learning model of RNA-protein binding

**DOI:** 10.1101/345140

**Authors:** Shreshth Gandhi, Leo J. Lee, Andrew Delong, David Duvenaud, Brendan J. Frey

## Abstract

**Motivation:** Determining RNA binding protein(RBP) binding specificity is crucial for understanding many cellular processes and genetic disorders. RBP binding is known to be affected by both the sequence and structure of RNAs. Deep learning can be used to learn generalizable representations of raw data and has improved state of the art in several fields such as image classification, speech recognition and even genomics. Previous work on RBP binding has either used shallow models that combine sequence and structure or deep models that use only the sequence. Here we combine both abilities by augmenting and refining the original Deepbind architecture to capture structural information and obtain significantly better performance.

**Results:** We propose two deep architectures, one a lightweight convolutional network for transcriptome wide inference and another a Long Short-Term Memory(LSTM) network that is suitable for small batches of data. We incorporate computationally predicted secondary structure features as input to our models and show its effectiveness in boosting prediction performance. Our models achieved significantly higher correlations on held out in-vitro test data compared to previous approaches, and generalise well to in-vivo CLIP-SEQ data achieving higher median AUCs than other approaches. We analysed the output from our model for VTS1 and CPO and provided intuition into its working. Our models confirmed known secondary structure preferences for some proteins as well as found new ones where secondary structure might play a role. We also demonstrated the strengths of our model compared to other approaches such as the ability to combine information from long distances along the input.

**Availability:** Software and models are available at https://github.com/shreshthgandhi/cDeepbind

## 1 Introduction

RNA binding proteins (RBPs) are crucial bio-molecules that play a key role in regulating many cellular processes, such as gene expression. RBPs fine tune gene expression by regulating various steps of pre-mRNAs processing, including splicing, editing and polyadenylation, and generate a large diversity of processed RNAs from the genome by regulating their maturation, stability, transport and degradation. For instance, HuR binds to target mRNA to enhance its stability and translation (de Silanes *et al.* (2004)) whereas, TIA-1 and TIAR suppress mRNA translation (Kim *et al.* (2011)). Due to their critical role in regulating post-transcriptional expression, mutations in RBPs or their binding sites can lead to many diseases including muscular atrophies and neurological disorders (Lukong *et al.* (2008); Musunuru (2003)). Since RBPs are involved in several stages of post-transcriptional regulation, understanding their binding preferences is crucial in understanding RNA processing, localisation and regulation.

RNA, being a single stranded molecule, folds onto itself, forming structures stabilised by hydrogen bonding between its bases. Due to the large diversity of structures of both RNA and RBPs the functional specificity of RBPs is complex and highly variable compared to DNA-binding proteins (Helder *et al.* (2016)). The structure of RNA can affect accessibility to target sites and affect binding of RBPs (Duss *et al.* (2014)). In addition to the complications due to structural context, RBPs such as some PUF proteins have been found to bind to multiple sites with variable spacing between them (Koh *et al.* (2009)). The protein PTB has been shown to contain four RNA binding domains (RBDs) all of which can bind to separate sites on RNA (Clerte and Hall (2006)). Multiple RBDs can contribute to overall binding affinity in a complicated non linear manner (Helder *et al.* (2016)).

Many RBPs are known to have a preference for both a specific sequence order and secondary structure of a portion of RNA (Hackermüller *et al.* (2005)). The shape of RNA which results from its folding is represented as base pairing between its nucleotides. There are several computational methods that predict the secondary structure of an RNA sequence from the order of its bases based on thermodynamic stability constraints (Steffen *et al.* (2005); Lorenz *et al.* (2011)). The structures can be represented as graphs (Janssen and Giegerich (2014)) or average probability vectors over the ensemble of all probable structures (Lorenz *et al.* (2011)). In addition, there are experimental methods that can probe RNA structures (Flynn *et al.* (2016)) but such data is scarce. It is commonly assumed that most proteins bind to accessible sites and few prefer a specific structural context (Li *et al.* (2014); Ray *et al.* (2009)). For instance, it is known that Vts1p is a yeast RBP that preferentially binds the sequence motif CNGG within RNA hairpins (Aviv *et al.* (2006)). Even though specific sequence preferences can be learnt from in-vitro experiments, a very small fraction (15%-40%) of specific RBP sequence motifs are occupied in-vivo (Taliaferro *et al.* (2016)). It has been shown that local secondary structure restricts access to a large subset of sequence motifs that would otherwise be bound (Taliaferro *et al.* (2016)).

Several in-vitro and in-vivo techniques have been developed to investigate RBP binding. High-throughput sequencing based experimental techniques such as CLIP-seq (Licatalosi *et al.* (2008); König *et al.* (2010); Hafner *et al.* (2010)), SELEX (Ellington and Szostak (1990); Stoltenburg *et al.* (2007)), RNAcompete (Ray *et al.* (2009, 2013)), RNA Bind-n-Seq (Lambert *et al.* (2014)) allow for the measurement of RBP binding affinities in a transcriptome-wide manner. Among the high-throughput techniques, the *in-vivo* methods HITS-CLIP, CLIP-seq and RIP-seq provide high quality test data for bench-marking binding models but owing to the high amount of noise and low resolution (Fu and Ares Jr (2014); Kishore *et al.* (2011)), training models on them directly is challenging. In-vivo experiments are further complicated by the presence of other RBPs and binding measurements could be the result of competition or complex formation between them. In-vitro experiments offer higher resolution, less noise and more accurate computational secondary structure prediction (Rouskin *et al.* (2014)). Hence, in this work we present results on training on in-vitro data, however we also note that our approach is agnostic to the data source and can be easily extended to train on in-vivo data as well. RNAcompete (Ray *et al.* (2009, 2013)) is a high-throughput in-vitro platform that measured the binding affinities (reflected as probe intensities) of over 200 RBPs to more than 240 000 probe sequences, each of length around 35 – 40 designed to cover every possible 9-mer at least 16 times. The large amount of data available for the experiments allows for training data-driven models such as deep neural networks and is the basis of the data used in this work. A new approach RNAcompete-S (Cook *et al.* (2017)) allows for querying a large diversity of RNA primary sequences and secondary structures and is an improvement over RNAcompete, which only allowed limited secondary structure representation and had probe sequences designed to be unstructured. The RNAcompete-S dataset is better suited to infer sequence-structure models since the probe sequences have stronger secondary structures. However, the data is available only for seven RBPs so far.

On the computational side, several methods have been developed to model RBP binding preferences including methods that rely on both sequence and secondary structure. The RNAcompete assay provided a comprehensive analysis of binding preferences of RBPs covering 205 genes and 24 different eukaryotes (Ray *et al.* (2013)). Several methods have been trained on data from RNAcompete including Deepbind (Alipanahi *et al.* (2015)), RNAcontext (Kazan *et al.* (2010)), and RCKOrenstein *et al.* (2016). Deep learning models learn distributed representations of data using multiple non-linear transformations and have shown promising results on several tasks where large amounts of data is available (LeCun *et al.* (2015)). Deepbind was the first approach to use deep neural networks to learn RBP binding preferences from RNAcompete data. It uses a convolutional neural network to model the mapping from sequence to binding intensity. However, Deepbind does not incorporate secondary structure as a feature and hence cannot model structural context. The specific architecture used for Deepbind also limits its ability to model complex interactions among multiple binding sites. RNAcontext incorporates structure in the form of structural context probability vectors that represented the ensemble of all possible structures. It learned a position weight matrix (PWM) to represent the sequence motif and global structure preferences to represent the preference of an RBP to bind to a given structural context. Since it only depends on a match between a PWM and local sequence, RNAcontext is position equi-variant and its predictions cannot depend on local secondary structure. RCK improved upon RNAcontext by using a k-mer model with local structure preferences. For each possible k-mer the RCK model learns a sequence score and structural context score. Due to its design that learns a weight for each possible k-mer, the number of weights for the model grows exponentially with *k*. The RCK method is thus limited to small values of *k* (*k* ≤ 6) and cannot model longer positional dependencies. RCK also averages the structural context along the span of each k-mer, and so cannot detect position-specific features. Graphprot is another approach that outperforms RNAcontext on sequence and structure specific binding(Maticzka *et al.* (2014)). It trains graph based support vectors by representing most likely structures as combinatorial graphs. As noted by the authors of (Orenstein *et al.* (2016)) Graphprot is prohibitively slow to run on the entire RNAcompete dataset as it has a runtime of about 7 days for each of the 255 proteins. Also it is unable to capture structural preferences from the RNAcompete data due to its unstructured nature.

To overcome the limitations of previous approaches, we augment the Deepbind model with structural information and refine its architecture to model more complex patterns of binding. We employ a Long Short Term Memory (LSTM) network and a convolutional neural network that allows our model to learn complex binding preferences. Our models can handle long range dependencies along RNA and identify structural preferences for RNA directly from data. Similar to the original Deepbind model, our model can be applied to any dataset where sequence and structure information is available. To benchmark our models we apply them to the RNAcompete dataset and significantly outperform state-of-the-art methods on in-vitro prediction. We demonstrate the ability of our models to capture complex interactions involving multiple binding sites. This work is based on SGs master’s thesis work made publicly available earlier at http://hdl.handle.net/1807/79240

## 2 Methods

### 2.1 Input and pre-processing

The models take as input a sequence *s* = (*s*_1_, …, *s_n_*) from the alphabet *A* = {*A, G, C, U*} and a structural annotation vector **r** = (*r*_1_, …, *r_n_*) where *r_i_* ∈ *R*^5^ and denotes the probability of the position being in the contexts (paired(P), hairpin loop(H), inner loop(I), multi-loop(M) or external region(E)). The structural context is computed using a variant of RNAplfold (Lorenz *et al.* (2011)) provided by the RNAcontext authors, that annotates the sequences into the aforementioned five structural profiles as opposed to the usual paired and unpaired profiles from RNAplfold. The sum of each structural context probability for a given *r_i_* sums to 1. The sequence input is encoded as a four-dimensional one-hot-encoding and is appended to the structural context vector to give a 9-dimensional input vector. The target scores are taken to be the RNAcompete probe intensities pre-processed to have the scores greater than the 99.95*^th^* percentile to be clamped at the value of the 99.95*^th^* percentile as done in Deepbind and RCK. Furthermore, the scores are normalised to have a mean of zero and a variance of one to ensure that consistent scales for learning rates and initial values of weights can be used without regard to individual intensity ranges for each protein

### 2.2 cDeepbind Model architecture

We develop two deep learning models for modelling RBP sequence and structure preferences. The models extend upon the original Deepbind model and can incorporate structural context, and are thus named Context-Deepbind or cDeepbind (Figure(1)). The cDeepbind-CNN model was designed with the aim to have a model with fast inference that could be applied to predict binding at each position on the transcriptome with single nucleotide resolution. The cDeepbind-RNN model can model more complex binding interactions but has slower inference. We use the mean-squared error between the predictions and target scores as our loss function for training. We also add an *l*_2_ penalty on the weights as a regularizer. For all our architectures we train an ensemble of identical models on the data with different random initializations and shuffled minibatches of data. Using an ensemble of models mitigates the problem of converging to local minima since by averaging the outputs we obtain a flatter minima which is known to generalise better(Kawaguchi *et al.* (2017)). We incorporated several techniques practised in the deep learning community to facilitate fast and efficient model training. We use Adam optimizer (Kingma and Ba (2014)) since it allows for stable training with recurrent architectures that have sparse gradients. We used batch size schedules (Smith *et al.* (2017)) which allows reduction in stochasticity in the gradients near convergence, by using increasing the batch size as training progresses. We implemented automatic hyperparameter search for our models which allows them to be applied to any new data source. We use GPU acceleration to allow us to train on the large amount of data in a rapid manner. We now describe both our architectures in detail.

**Fig. 1.**
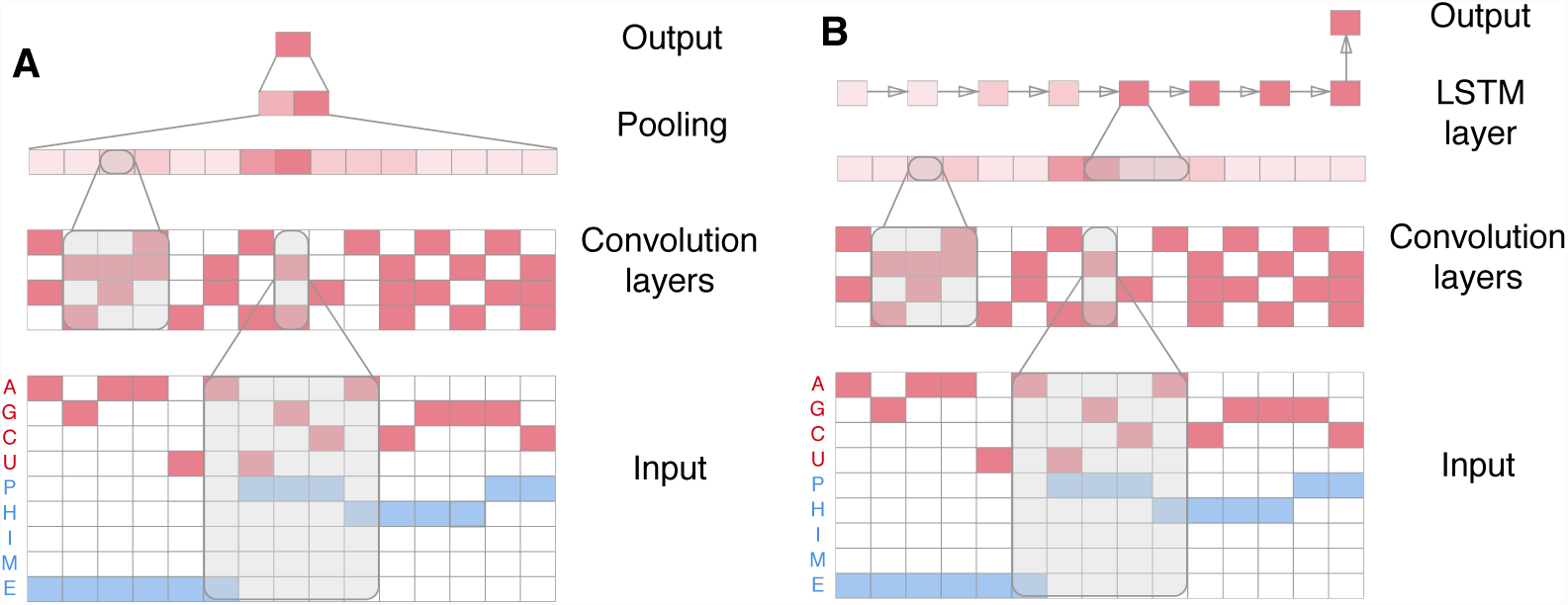
(a) The cDeepbind-CNN architecture uses two convolution layers followed by global max and average pooling. The values of the feature map in the second convolution layer **q** is a summary of local information within its receptive field. Thus, the value in *q_k_* depends on (*s_k_*, …, *s*_*k*+*w*_), where *w* is the receptive field of units in **q**. The pooling then combines these features along the sequence length to produce the final prediction (b) The cDeepbind-RNN architecture uses an LSTM layer to process the feature map of the convolution layers to produce a summary vector **o**. Since each sequential output in the LSTM is influenced by past inputs in the sequence *o_k_* depends on (*s*_0_*, …, s_k_*_+*w*_), where *w* is the receptive field of units in **q**

#### 2.2.1 cDeepbind-CNN

The cDeepbind-CNN model uses a fully convolutional architecture to predict binding intensity. Convolutional Neural Networks(CNNs) are useful where local groups of data are highly correlated and form distinct patterns or motifs. For instance, in image models, they preserve local correlations in the data, such as edges, curves, and contours. Combinations of lower level features give rise to more complex shapes such as faces. Analogously, for genomic sequences convolutions can detect motifs along the sequence and combine them to create a representation for the entire input. Here we use two convolution layers with ReLU hidden units to transform the input. The choice of ReLU activation allows efficient backpropagation of gradients since it does not saturate. We do not use any sub-sampling in the convolution layers to retain single nucleotide resolution at the final convolution output. We then perform max and average pooling across the entire sequence length to get a two-dimensional score which is then weighted to produce the final binding score. The global pooling layers are used to generate a summary score for the entire input using the representations learned at the lower layers. Using global pooling also imparts the model flexibility in the length of the input. Since there is no sub-sampling or fully connected layers, the sequence order is retained in the final convolution output. Thus, from the model, it is possible to produce a binding track that shows the binding scores for each position in the sequence.

#### 2.2.2 cDeepbind-RNN

The cDeepbind-RNN model uses a convolutional LSTM architecture. Recurrent neural networks (RNNs) are networks for processing sequential data. They do this by sharing parameters across time steps and thus are able to share useful representations between them. RNNs process the input sequence one element at a time, maintaining a state vector that contains a summary of all observations seen so far. The input at each time step is used to update the hidden state. The output of the time step is produced by a non-linear combination of the input at the given time step and the state from the previous time step. A Long Short Term Memory (LSTM) network is a recurrent network that can handle long dependencies along the data without losing information through gradient backpropagation. We use an LSTM since it can efficiently capture non-linear interactions between different input positions such as combinations of multiple binding sites, specific flanking structural context etc, by storing past context in its internal hidden state. In our architecture, the convolution layers recognise local motifs along the sequence and pass that as input to the LSTM. The LSTM then processes the convolution output sequentially and produces the final binding prediction at its last time step.

### 2.3 Hyperparameters

Deep learning models are known to be sensitive to hyperparameters. Random sampling of hyperparameters generally performs better than grid search or manual tuning (Bergstra and Bengio (2012)). We perform hyperparameter search by randomly sampling 5 hyperparameters for each model run and choose the one with the lowest validation cost on three-fold cross validation. We sample the filter widths for the convolution layers in the range 8 – 16, the number of filters in the range 8 – 24, the number of hidden units for the LSTM in the range 10 – 30. We use random normal initialization for the weights and initialize the biases to a small positive value.

### 2.4 Training pipeline and hyperparameter tuning

The model training pipeline consists of automatic hyperparameter training using cross validation followed training an ensemble of models with the best hyperparameter from the previous step on the entire training set. We use an ensemble of five models as our final predictive model by averaging the predictions across the ensemble. We find that the most important hyperparameters for the model were the learning rate, rate of weight decay, number of LSTM units and variance of the random normal initializer for the weights. We use batch-size scaling which is a new alternative to learning rate decay, to accelerate training. We start training with a minibatch size of 500 and double the batch size after every 5 epochs. We found during cross-validation that loss saturates after 15 epochs and we train all our models for 15 epochs. The code for the model was written in Tensorflow (Abadi *et al.* (2016)) and the models were run on NVIDIA Tesla K80 GPUs. The training time including hyperparameter search for a single protein on one GPU was around 30 minutes for cDeepbind-RNN and 10 minutes for cDeepbind-CNN. We use parallel training across 15 GPUs to train the entire list of 255 proteins in 3 hours for the cDeepbind-CNN and 8 hours for cDeepbind-RNN.

## 3 Results

We compared our models with other methods such as RCK, Deepbind and RNAcontext. We observed that our model outperforms all other approaches and can learn the relevant sequence and structural preferences for the proteins under study.

### 3.1 cDeepbind is more accurate at in vitro binding prediction than state of the art

We compared the performance of our model on predicting probe intensities on the RNAcompete dataset (Ray *et al.* (2013)) across 244 experiments. We find that even though the probes in this experiment were designed to be unstructured we can still learn some structural preferences from it, which was discovered by the authors of RCK as well. The data consists of a training set of sequences (set A) and a held out test set (set B). The models are trained on set A and the performance measure is taken as the Pearson correlation of predicted intensities with the probe intensities on the test set. We used the results published in the RCK study (Orenstein *et al.* (2016)) for comparison against other methods. As illustrated in Figure (1), both our models significantly outperform all other methods on in-vitro binding prediction. We obtained an average Pearson correlation of 0.594 and 0.5336 for our RNN and CNN models respectively. We outperforms the state of the art method RCK, which achieves an average correlation of 0.461 and the original Deepbind model which obtains an average correlation of 0.435 (P-values using Wilcoxon signed-rank test in Figure (1,**B**,**C**)).

We found that the average relative improvement in correlation over RCK was 33.21% and 18.15% over the 244 experiments for the RNN and CNN models respectively. We believe the improvement in performance results from the model being able to recognise longer patterns, complex interactions, and interplay across sequence and structure components of the input.

### 3.2 cDeepbind is sensitive to secondary structure

To evaluate the impact of secondary structure in the improvement of predictions we trained the cDeepbind-RNN model without secondary structure as a feature. Both sets of models were trained with an LSTM output layer and the same hyperparameter search space for the sizes of the hidden units. We found that the model that has access to secondary structure features performs better with an average Pearson correlation of 0.594 versus 0.528 for the model that does not use secondary structure features (P-value=3.91 × 10^−39^, Wilcoxon signed rank test). We find that the relative improvement is greatest in some proteins that have a known secondary structure preference such as RBFOX1. We also discovered that several other proteins had a big improvement in predictions upon including secondary structure, such as SF1, PUM, MSI, and others.

### 3.3 cDeepbind discovers known structural preferences for VTS1p

To visualise the binding preferences learned by the cDeepbind-RNN model we looked at some sequences that had the highest experimentally observed binding intensities for the protein and visualised the gradient of the loss function with respect to the input. The gradient acts as a sensitivity map that tells us what the model is paying attention to. It should be noted here that the input to the model are one-hot vectors which are a continuous relaxation of the original discrete sequences. Even though the model treats the input space as continuous, it is useful to think of the direction of the gradient at a given position being an indicator of a nucleotide switch at that position resulting in either an increase or decrease in the overall loss. We illustrate the sensitivity of the model to different parts of the input using mutation maps (Figure (3,**A**,**B**)), which were first developed in the original Deepbind study. The mutation map is a colour map with four rows corresponding to the bases A,G,C,U in the input from top to bottom. The cells are coloured on a spectrum from blue to red with white meaning zero gradient (Figure (3, **C**)). A negative gradient indicates a reduction in the loss which corresponds to a prediction closer to the target score. Hence, the red cells correspond to positions useful for binding whereas blue cells correspond to cells that would prefer a different nucleotide. The size of the letters on top of the mutation map corresponds to the relative magnitude of the gradient at that position, with positions more relevant to the final predictions having a greater size. We also overlay the gradient on to the predicted secondary structure computed using RNAfold (Hofacker (2003)) as shown in Figure (3, **D**). Our model confirms the known binding preference of VTS1p to bind to CNGG motifs within hairpin loops. The mutation map (Figure (3, **A**)) shows that when the model encounters the CNGG motif(highlighted in the red rectangle in Figure (3, **A**)), it is sensitive to the C and two Gs, with all other nucleotides in their positions predicting a decrease in score and the model does not care about the nucleotide sandwiched between the C and G since it shows almost a zero gradient in that cell. The model also gives a high weight to the gradient at the structure component of the input around the CNNG motif being in a hairpin state and the flanking region being paired, as seen in Figure (3, **B**), which corroborates the known binding preference of VTS1.

**Fig. 2.**
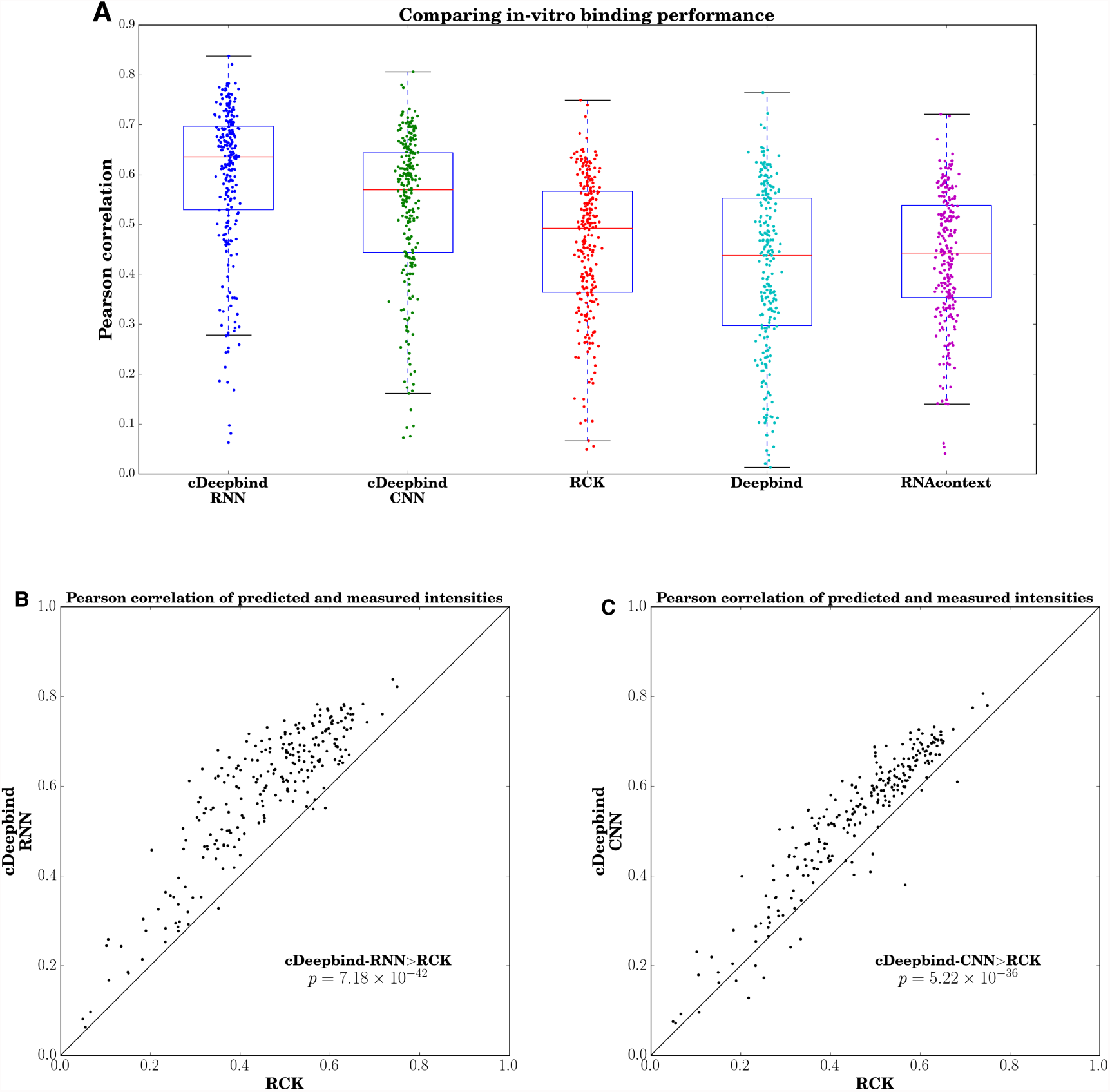
(A) Box plot comparing the performance of extant methods on in-vitro binding prediction on the RNAcompete dataset. All models were trained on Set A sequences and tested on Set B. Dots represent Pearson correlation on the test set for 244 experiments. (B) Scatter plot comparing cDeepbind-RNN to state-of-the-art method RCK. (C) Scatter plot comparing cDeepbind-CNN to RCK.

**Fig. 3.**
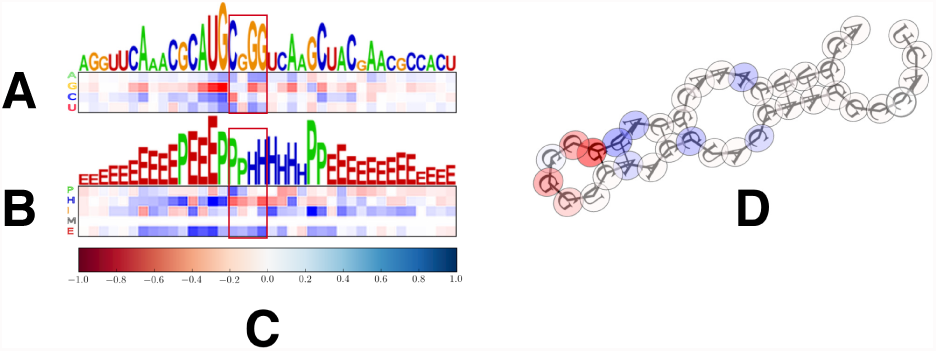
Mutation map for the sequence(A) and structure(B) components of the input, and Minimum Free Energy (MFE) structure (D) for a sequence that binds VTS1 with high affinity using the cDeepbind-RNN model. The mutation map is along the length of the input with the gradient normalised for each position. The cells are coloured according to the normalised gradient for each possible input according to the colour-bar in (C)

### 3.4 Comparing CNN and RNN models for CPO

We take a sequence from the test set of the CPO model that has normalised target score of 26.90. The cDeepbind-RNN model assigns a prediction of 23.91, whereas the cDeepbind-CNN model assigns a score of 4.1. CPO is a protein that is known to bind to the *GCAC* motif (Ray *et al.* (2013)). As illustrated in Figure (4), both the models are sensitive to the *CAC* motifs within the sequence(highlighted within red rectangles in Figure (4,**A**,**B**)), with the gradient suggesting that any alterations in those positions in the sequence would lead to a greater mean squared error with the target score. The gradient also suggests that the models would have preferred a *G* instead of a *U* preceding the *CAC* at the far right of the sequence which agrees with the literature motif for CPO. We also see here that the CNN model focuses on just one motif and does not take into account the combined effect of the multiple motifs in the sequence (Figure (4, **B**))

**Fig. 4.**
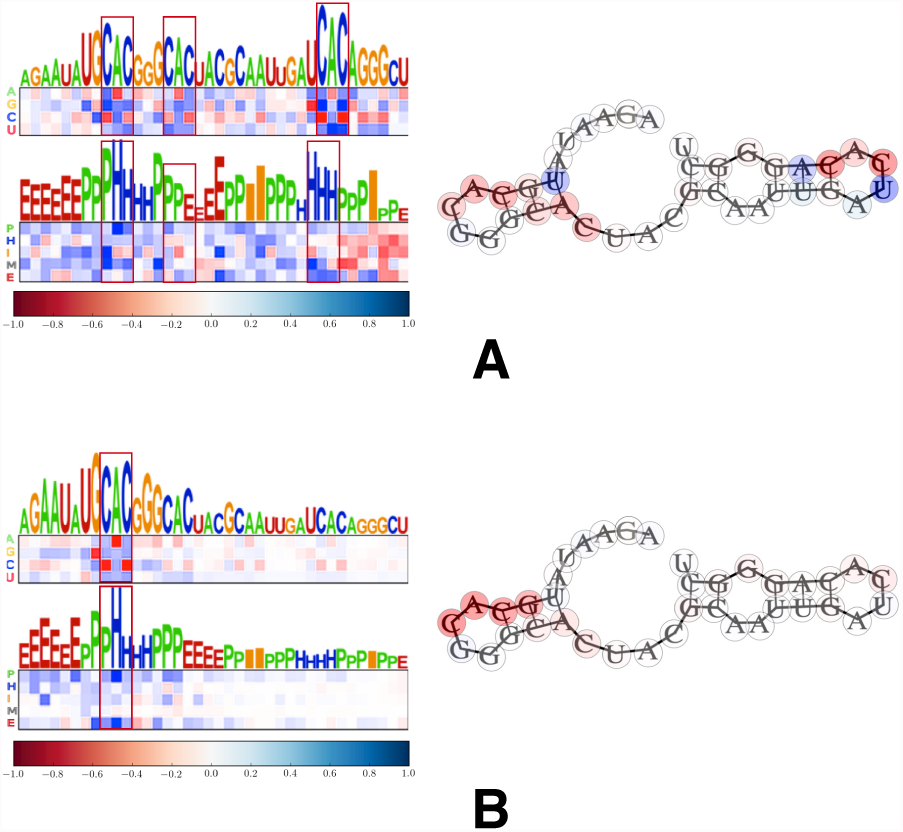
Mutation map and MFE structure for a sequence that binds CPO with high affinity using the (A) cDeepbind-RNN model and (B) cDeepbind-CNN model.

The RNN however, can handle non-additive effects of multiple motifs in a better way due to its architecture and is able to make a prediction closer to the target score. The RNN model seems to suggest that the CPO protein might prefer multiple *CAC* motifs in conjunction, for binding. This is corroborated by recent work where the binding specificity of the protein RBPMS was investigated (Teplova *et al.* (2016)). The RRM domain of RBPMS was reported to bind to a pair of tandem *CAC* motifs spaced by a variable nucleotide window. It was also reported that the majority of the residues of the RNA recognition motif (RRM) domain of RBPMS were strictly conserved in the corresponding RRM of CPO. Thus it is reasonable to assume similar binding patterns for RBPMS and CPO. This example illustrates the ability of the RNN model to combine information along the sequence in a non-additive way and its advantage over the CNN in handling such dependencies.

### 3.5 Comparison on in vivo binding data

We evaluated our model on 23 pairs of RNAcompete and CLIP experiments that covered 10 proteins. The CLIP experiments were taken from the GraphProt study (Maticzka *et al.* (2014)). The CLIP datasets contained in vivo binding sites collected from CLIP experiments. Control sequences were extracted from unbound regions of the same bound transcripts. The dataset contained flanking sequences of length 150 nucleotides on both sides of both bound and control sequences. We used the sequences including flanks for secondary structure prediction with RNAplfold while only the sequence and predicted probabilities were used for testing. Since the RCK paper bench-marked models trained on the entire set of sequences including the test set, we retrained models for this comparison on the larger set of sequences. The performance values for RCK, RNAcontext, and Deepbind were taken from the RCK study.

We compared the cDeepbind-CNN and cDeepbind-RNN models against other methods (Figure (5)). The AUC for a protein was taken as the average AUC for all pairs of RNAcompete-CLIP experiments that covered the proteins. To compare against other methods we consider the median of the average AUCs for the 10 proteins. We find that our both our CNN and RNN models obtain higher AUCs than other methods achieving median AUCs of 0.840 and 0.805 respectively against 0.803 for RCK and 0.790 for Deepbind. The p-values between cDeepbind-CNN and the next best performing method, RCK, was 0.029, whereas the p-value between cDeepbind-RNN and RCK was 0.057 (Wilcoxon signed-rank test). We believe we do not see as big of a boost in performance in-vivo as compared to in-vitro because of the noisy nature of the data and inaccuracy of secondary structure prediction in-vivo. In addition, there are very few proteins in the in-vivo evaluation set to quantify improvements accurately.

**Fig. 5.**
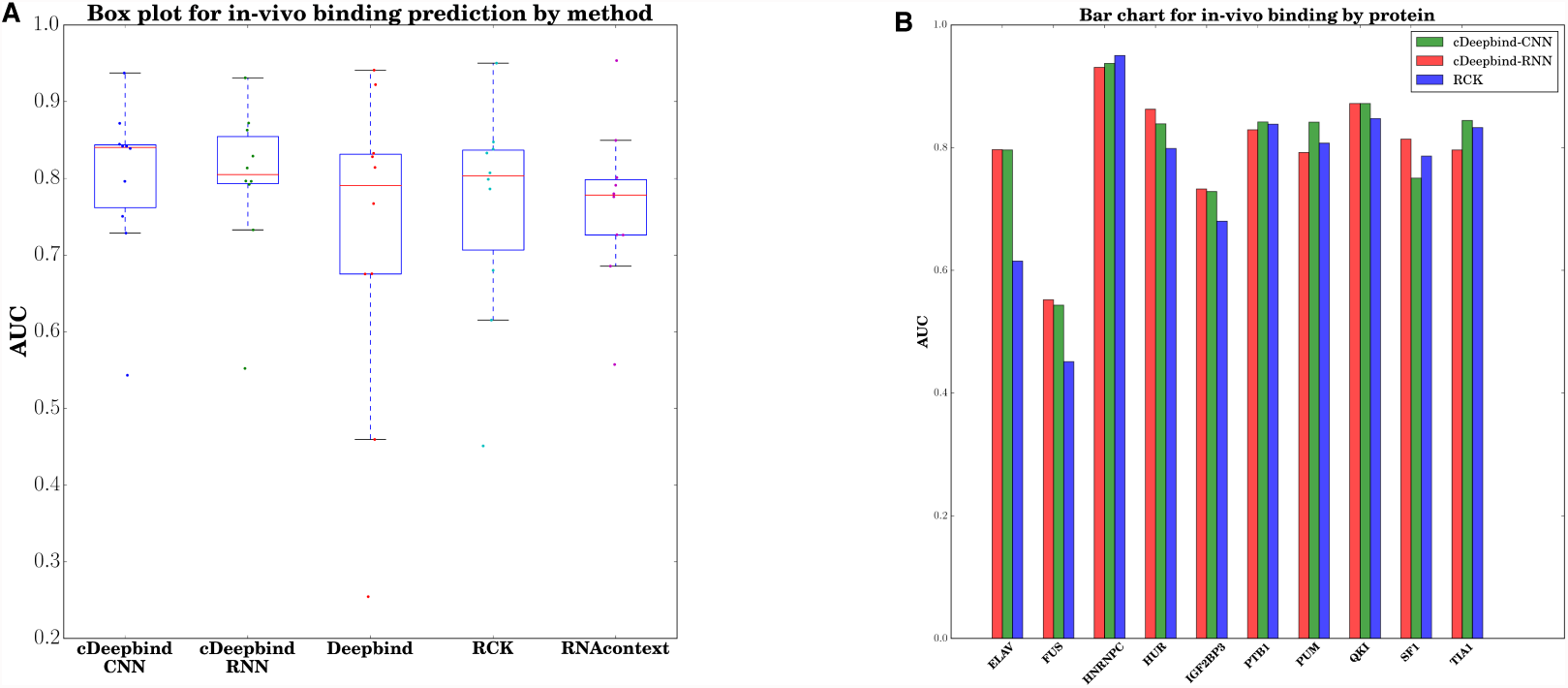
(A) Box plot comparing the performance of our models with previous approaches. Each dot represents average AUC across different CLIP experiments for a protein. (B) Bar plot illustrating the average AUCs for each protein for cDeepbind-RNN, cDeepbind-CNN and RCK. We obtain an improvement in median AUC with our models.

## 4 Conclusions

We have presented a deep learning approach to model RBP binding preferences that incorporates both sequence and secondary structure information. Our approach provides an improvement over k-mer models by using distributed representations in the form of convolutional filters to model the binding preferences. Our models can learn preferences for long motifs in the input. We show that our CNN model can produce binding tracks from the input and our RNN model can store past context in its internal state to make better predictions. Our models significantly outperform previous methods on in-vitro evaluation based on Pearson correlation on held out test data. On in-vivo evaluation, we achieve improvement in median AUC on both of our models, albeit the improvements are not very significant. We believe in-vivo data for more proteins that overlap with the set of proteins available in RNAcompete, would demonstrate the improvement in performance more significantly.

Our models show a large gap in performance with and without secondary structure information for some proteins that are known to be sensitive to secondary structure such as VTS1p and RBFOX1. In addition, we discover such difference for other proteins such as SF1, PUM and MSI which do not have known secondary structure preferences.

Even though we applied our model to the RNAcompete in-vitro data, our approach is general and the model can learn relevant features from any data source including in-vivo CLIP datasets. It would be useful in the future to train and validate on RNA-bind-n-seq and RNAcompete-S. Future directions for this work could include training on in-vivo data, training a multitask prediction model for similar proteins, and generative modelling of binding sequences.

## Acknowledgements

We would like to thank Quaid Morris for his helpful comments and suggestions.

